# Does juvenile Baltic sturgeon (*Acipenser oxyrinchus*) smell the enemy?

**DOI:** 10.1101/476952

**Authors:** Maria Cámara Ruiz, Carlos Espírito Santo, Alfons Mai, Jorn Gessner, Sven Wuertz

## Abstract

Atlantic sturgeon (*Acipenser oxyrinchus*), also known as Baltic sturgeon, is considered extinct in German waters. Fish-rearing for conservation purposes still relies on classical hatchery technology producing fish not well suited for facing life in the wild, lacking behavioural skills such as foraging or anti-predation. Predation is hence a major source of mortality in newly stocked individuals. The aim of this study was to evaluate if naïve Baltic sturgeon juveniles were able to smell and recognize a common predator, sander (*Sander lucioperca*). Over a period of 30 days, three tanks from each group of Baltic sturgeon were attached to a rearing tank with sander (sander unit) and, as a control, carp (carp unit). Morphology of the dorsal scutes, distribution within the tank, stress (glucose, lactate and cortisol) and gene expression of brain plasticity and cognition were studied in comparison to the control group (carp unit). No significant differences were observed in any of the parameters measured. Thus, we conclude that naïve Baltic sturgeon is not able to innately recognize potential predators and future studies should focus on implementing predator odour together with chemical alarm substances.

## 1. Introduction

Sturgeons (Acipenseridae) have experienced a drastic decline due to several reasons including overfishing, habitat destruction and pollution over the last decades. Nowadays, sturgeons are among the most endangered fish species worldwide (IUCN, 2011). Baltic sturgeon (*Acipenser oxyrinchus)* has been indigenous to the Baltic region, but it is currently considered extinct (Gessner et al., 2006; Langhans et al., 2016). In order to recover Baltic sturgeon populations, restoration programs have been established to reestablish the species in its former distribution range by releases of juveniles with hatchery origin in an attempt to build up self-sustaining populations (Gessner et al., 2011).

Fish-rearing for conservation purposes still relies on classical aquaculture hatchery techniques that focus on growth, survival and reproduction within the hatchery, which are known to have shortcomings with regard to the behavioural skills needed in the wild (Ferno & Jarvi, 1998; Olla et al.,1998). One of the pitfalls of these methods is the high mortality of newly stocked individuals (Brown & Smith, 1998; Suboski & Templeton, 1989). In juvenile fishes, predation is a major source of the post-stocking mortality following release (Brown & Day, 2002; Brown & Laland, 2001).

Prey animals can determine risk by using a variety of visual, chemical and mechanical cues (lateral line). Regarding chemical cues, fishes heavily rely on chemosensory information, specifically semiochemicals. There are three classes of semiochemicals: kairomones, disturbance cues and damage-released alarm cues. Kairomones are cues emitted by one species and are detected by another, for instance, the scent of predators detected by prey. Kairomones are adaptively favourable to the receiver and help prey to detect and avoid potential predators (Ferrari et al., 2010).

The possibility of training captive-bred animals in predator avoidance prior release into the wild has received attention in the conservation context (Olla et al., 1994; Brown & Laland, 2001; Griffin et al., 2000; Wallace, 2000). Exposure to various predator-stimuli prior to reintroductions has already been used with mammalian and avian prey (Griffin et al., 2000), but has received much less attention in fishes (Brown & Day, 2002). Antipredator behaviour is often assumed to be strongly defined by genetic components, but fishes can be very flexible in adjusting their responses (Kelley & Magurran, 2003). Studies have shown that the appropriate stimulus is able to improve the avoidance responses of fishes (Berejikian, 1995; Brown & Smith, 1998; Mirza & Chivers, 2000)

In this study, the objective is to determine if juvenile Baltic sturgeon is able to recognize a common predator, sander (*Sander lucioperca*) by smell. Therefore, sturgeon juveniles were kept in the rearing water of sander for 30 days. Distribution within the tank, stress (glucose, lactate and cortisol), morphology of the dorsal scutes and gene expression of brain plasticity and cognition markers were studied in comparison to a control group that was reared in water used to rear carp (Fig. 1).

**Figure 1.**
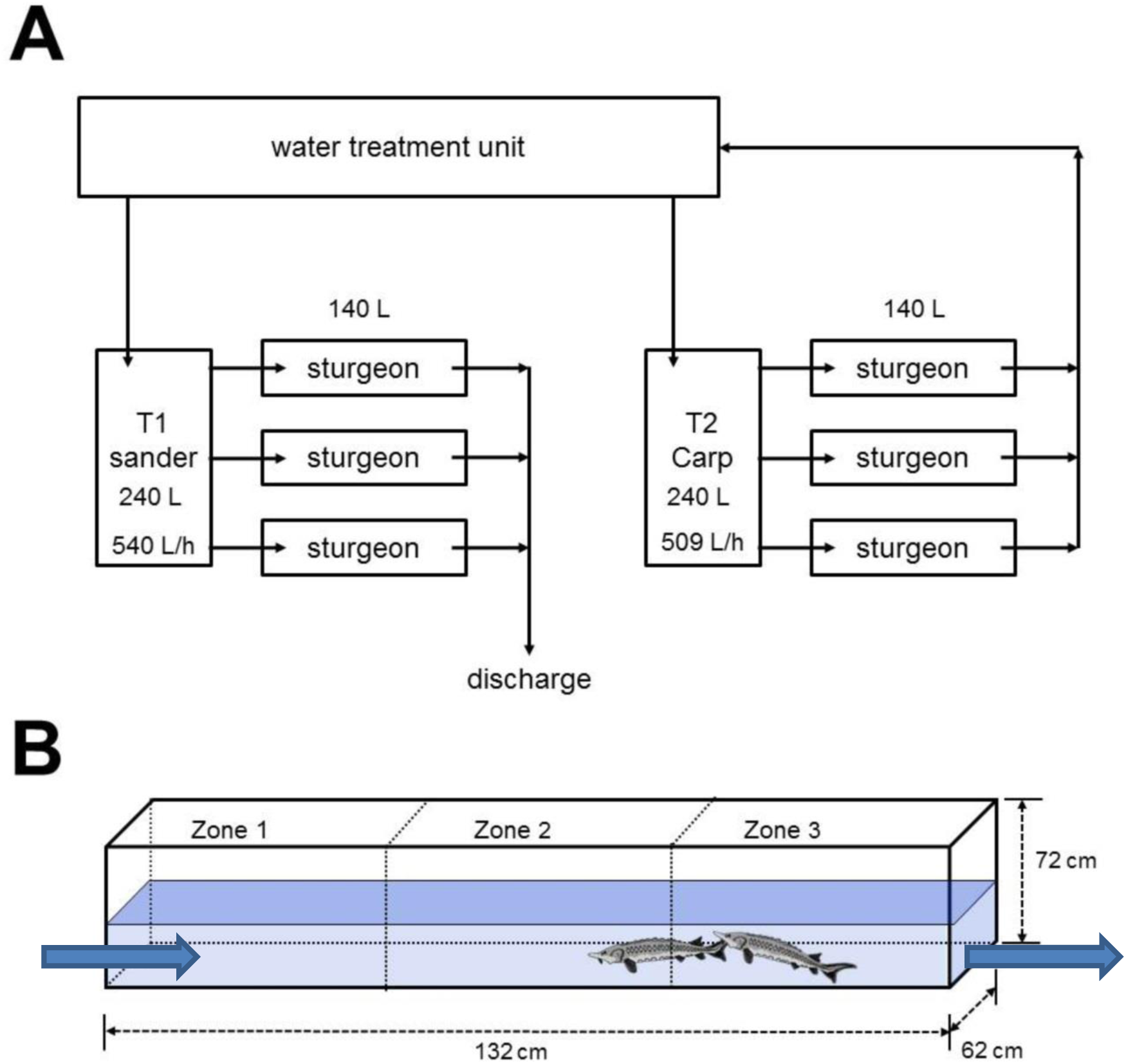
Experimental design. 1A. Three tanks from each group were attached to a rearing tank with sander (sander unit) or, as a control, carp (carp unit). 1B. Tank distribution. Different zones were established in order to observe the distribution towards the inlet.

## 2. Materials and Methods

### 2.1. Experimental design

A total of 120 juvenile Baltic sturgeon, *Acipenser oxyrinchus*, (23.5 ± 2 cm; 39.9 ± 8.8 g) were randomly distributed to 6 tanks (132 cm* 62 cm* 72 cm; V = 140 l). Three tanks of each group was attached to a rearing tank with sander (sander unit) or, as a control, carp (carp unit). The sander units were maintained in flow through with water of the municipal water supply, the carp rearing water was recirculated through a filter unit. The tanks were stocked at 4 kg/m^3^ corresponding to 10 sander and 13 carp per tank (Fig. 1A). Water turnover was 540 l/h and 509 l/h in the sander (T1) and carp tank (T2) respectively. In the sturgeon tanks, water turnover was 172 ± 7.2 l/h. Fish were kept at a temperature of 19.8± 0.4 under a natural photoperiod. Water parameters (T, O_2_ 9.1 mg/L) were monitored daily, nitrite (< 0.05 mg/L) and TAN (< 0.09 mg/L) were determined every 3 days. All fish (sturgeon, sander and carp) were allowed to acclimate for two weeks prior to the start of the experiment.

### 2.2. Sampling procedure

Sturgeon juveniles were sampled after 30 days of experiment. Blood samples were collected from the caudal vein with a syringe. Serum was collected after centrifugation (10,000 rpm, 5 min, 4° C) and stored at −20° C until use. Scute morphology was assessed by photographs. The increase in length of the protrusions was determined before and after the experiment with the ImageJ program (https://imagej.nih.gov/ij/). At the end of the experiment, the fish were killed by an overdose of MS222 and the brain was removed surgically. Sturgeon brains were dissected and divided into three parts representing the three main brain regions (forebrain, midbrain and hindbrain). Samples were stored in RNA later at −80 °C until gene expression analysis.

All experiments were in compliance with EU Directive 2010/63/EU and approved by the national authorities (G0305/15, Landesamt für Gesundheit und Soziales, Berlin, Germany).

### 2.3. Tank distribution

Distribution of sturgeon in the rearing tank was determined from 7 photos taken every 5 min at 1 day, 5 days, 10 days, 20 days and 30 days of the experimental trial. In order to assess the distribution, the tank was divided into 3 zones (up-, mid-, downstream section of the through). The juveniles recorded in each zone expressed as percentage of the total number of fish in the tank was recorded.

### 2.4. Scute morphology

To determine the morphology of the scutes, three scutes anterior to the dorsal fin were analyzed using ImageJ (https://imagej.nih.gov/ij). The measurements included diagonal (distance from the basis to the tip of the respective scute) and basal distance. The measurements were calibrated to a scale bar. All measurements were carried out in triplicate.

### 2.5. Blood parameters (glucose, lactate and cortisol)

For the determination of serum parameters, blood was sampled from the caudal vein with a heparinized syringe. After centrifugation at 5000 g for 5 min, cell-free plasma was immediately shock frozen and stored at −80 °C until further processing.

Glucose in the plasma of Baltic sturgeon was measured with a glucose colorimetric GOD-PAP kit (Greiner). In order to initiate the reaction, 5 μl of serum (1:5 dilution) from Baltic sturgeon were mixed with 250 μl of reagent. The mixture was incubated for 10 min at 25° C. Afterwards, the absorbance was measured at 500 nm and concentrations were calculated from a dilution series (0.0625-1 mg/ml).

Plasma lactate in the serum of Baltic sturgeon was measured with a lactate colorimetric LOD-PAP kit (Greiner). In brief, 5 μl of serum (1:2 dilution) from Baltic sturgeon was mixed with 250 μl of reagent. The mixture was incubated for 10 min at 25° C. Afterwards, the absorbance was measured at 500 nm and concentrations were calculated from a dilution series (0.0375-0.3 mg/ml).

Cortisol was determined using an ELISA kit (IBL, Germany). In brief, 100 μL plasma was extracted by vigorously shaking for 3 min with 1.9 mL ethanol in 5 mL glass vials. The organic phase was transferred to a new vial and the extraction was consecutively repeated twice as described above. The three fractions were pooled and allowed to evaporate under a constant nitrogen stream. For analysis, the remaining steroid fraction was redissolved in assay buffer (IBL, Germany). Corstisol concentrations were determined in duplicate at 450 nm with an Infinite M200 microplate reader (Tecan, Germany) and calculated from a standard dilution series.

### 2.6. Gene expression

Total RNA was extracted with TRIzol as described by Reiser et al., (2011), including a DNase I digestion. Total RNA concentration and purity were determined in duplicates with a Nanodrop^®^ ND-1000 UV–Vis spectrophotometer. Purity was validated by the ratio of the absorbance at 260 and 280 nm (A260/280) ranging between 1.8 to 2.0. Moreover, integrity of the total RNA was checked by gel electrophoresis and, in 10% of all samples, on a RNA 6000 Nano chip with an Agilent 2100 Bioanalyzer. To eliminate potential DNA contamination, DNAse I digestion was performed in all samples prior to transcription. Next, mRNA was transcribed with MMLV Affinity reverse transcriptase (Agilent, 200 Units/μl) according to the manufacturer’s instruction. In 10% of the samples, the enzyme was substituted by pure H_2_O, serving as a control (-RT) to monitor DNA contamination.

Species-specific primers targeting elongation factor 1α (*ef1a*), brain-derived neurotrophic factor (*bdnf*), neurogenic differentiation factor (*neurod1*) and proliferating cell nuclear antigen (*pcna*) were designed using the sequence information available. Specificity of the assays was confirmed by direct sequencing (SeqLab, Germany). Real-time PCR was carried out with Mx3005p qPCR Cycler (Stratagene), monitoring specificity by melting curve analysis.

For the PCR reaction, 2 μL of the diluted samples (40 ng/μL) were used as template in 20 μL PCR mix [SYBR-Green I (Invitrogen), 200 μM of each dNTPs (Qbiogene), 3 mM MgCl 2 and 1 U Invitrogen Platinum Taq polymerase]. PCR conditions comprised an initial denaturation at 96 °C for 3 min, followed by 40 cycles of denaturation at 96 °C for 30 s, primer annealing (for Ta, see Table 1) for 30 s and elongation at 72 °C for 30 s. PCR efficiencies were determined experimentally with a dilution series of a calibrator corresponding to 200 ng/μl. PCR assays for all individual samples were run in duplicate. Expression of target genes were calculated by the comparative CT method (ΔΔCT) according to (Pfaffl, 2001), correcting for the assay efficiencies and normalizing to *ef1a* as a housekeeping gene. Expression data are presented as fold increase of the respective calibrator.

**Table 1.**
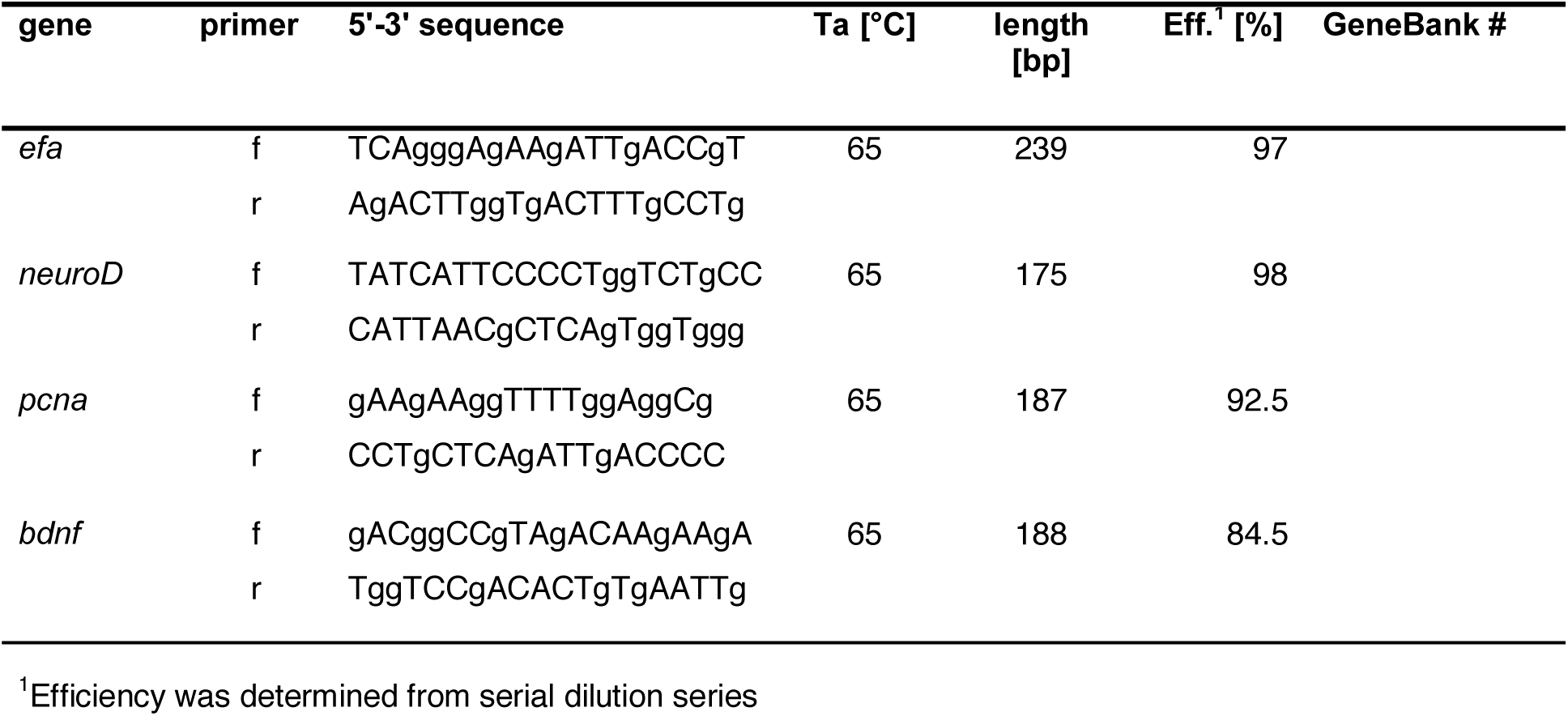
Specifications of qPCR assays including primer sequences, annealing temperature (Ta), amplicon length [bp], PCR efficiency (Eff) and NCBI accession number of the respective housekeeping (ef) and target genes: ef-elongation factor 1 a, neurod1-neurogenic differentiator factor, bdnf - brain-derived neurotrophic factor, pcna-proliferating cell nuclear antigen-

### 2.7. Data analysis and statistical methods

Data are presented as mean ± standard deviation (SD). Prior to statistical analyses, all data were tested for normality of distribution using the Kolmogorov-Smirnov/ Shapiro Wilk test and for homogeneity using Levene’s test. Data on the behavior were scored and analyzed using Dunn’s multiple comparison test. RNA expression, lactate, glucose and cortisol were analysed with a parametric student T-test or non-parametric Mann-Whitney test. The level of significance used was P ≤ 0.05. All statistical analyses were performed with GraphPad Prism statistical program.

## 3. Results

### 3.1. Tank distribution

The distribution of the fish in the tanks did not reveal significant differences for the percentage of Baltic sturgeon recorded in each zone (Fig 2) over time. Approximately, 25-35% of sturgeon stayed in each of the three zones regardless of the source of the inflow water. No significant differences were observed between dates or treatment observed (Dunn’s, P ≤ 0.05).

**Figure 2.**
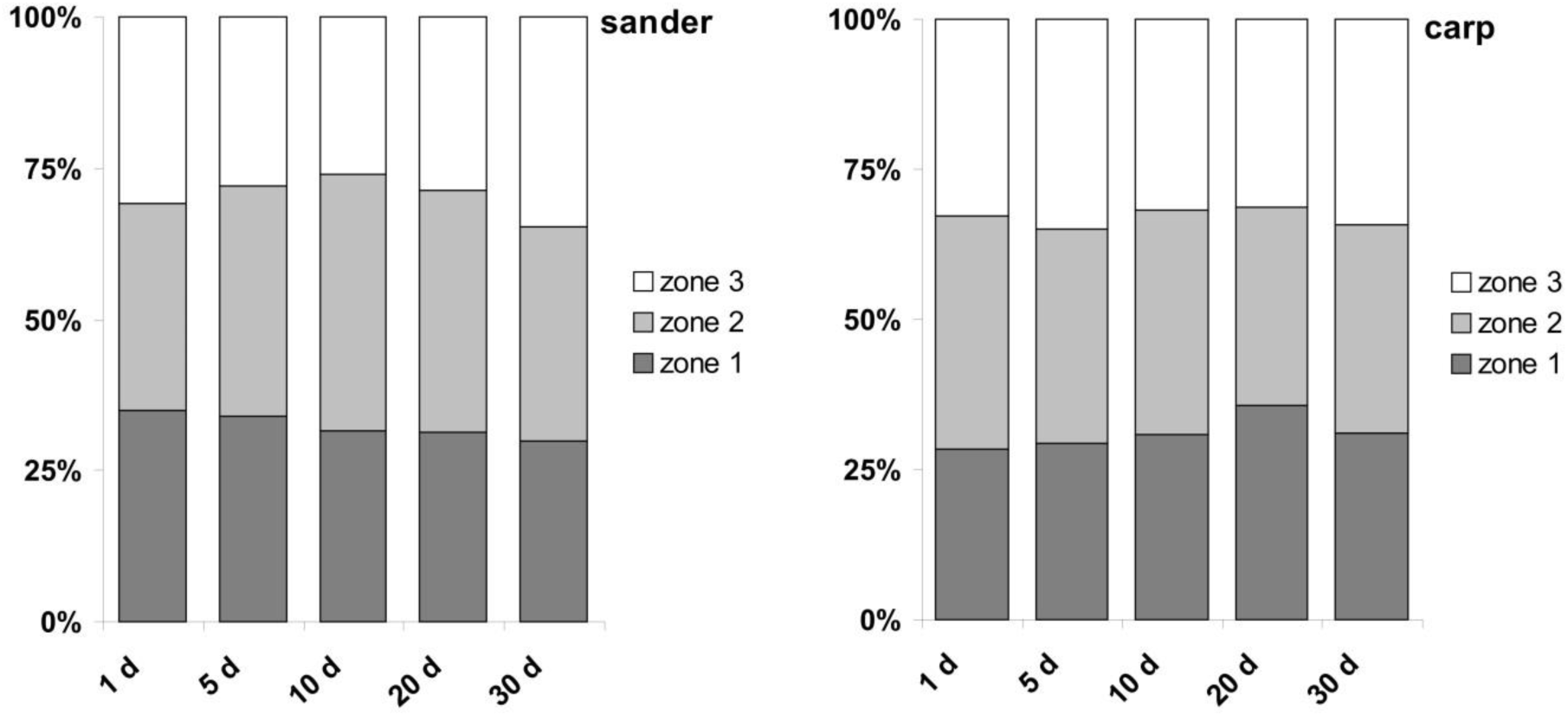
**Tank distribution.** Percentage of Baltic sturgeon (*A. oxyrinchus*) in each zone of rearing troughs during the exposition to water from Sander (left) or carp (right) rearing systems. No significant differences over time (Dunn’s, p ≤ 0.05) or between treatments (Mann-Whitney test, p ≤ 0.05) were observed.

### 3.2. Scute morphology

Diagonal and basis length of the scutes were correlated (Fig 3). There were no significant differences of the coefficient between the sander and the carp group observed.

**Figure 3.**
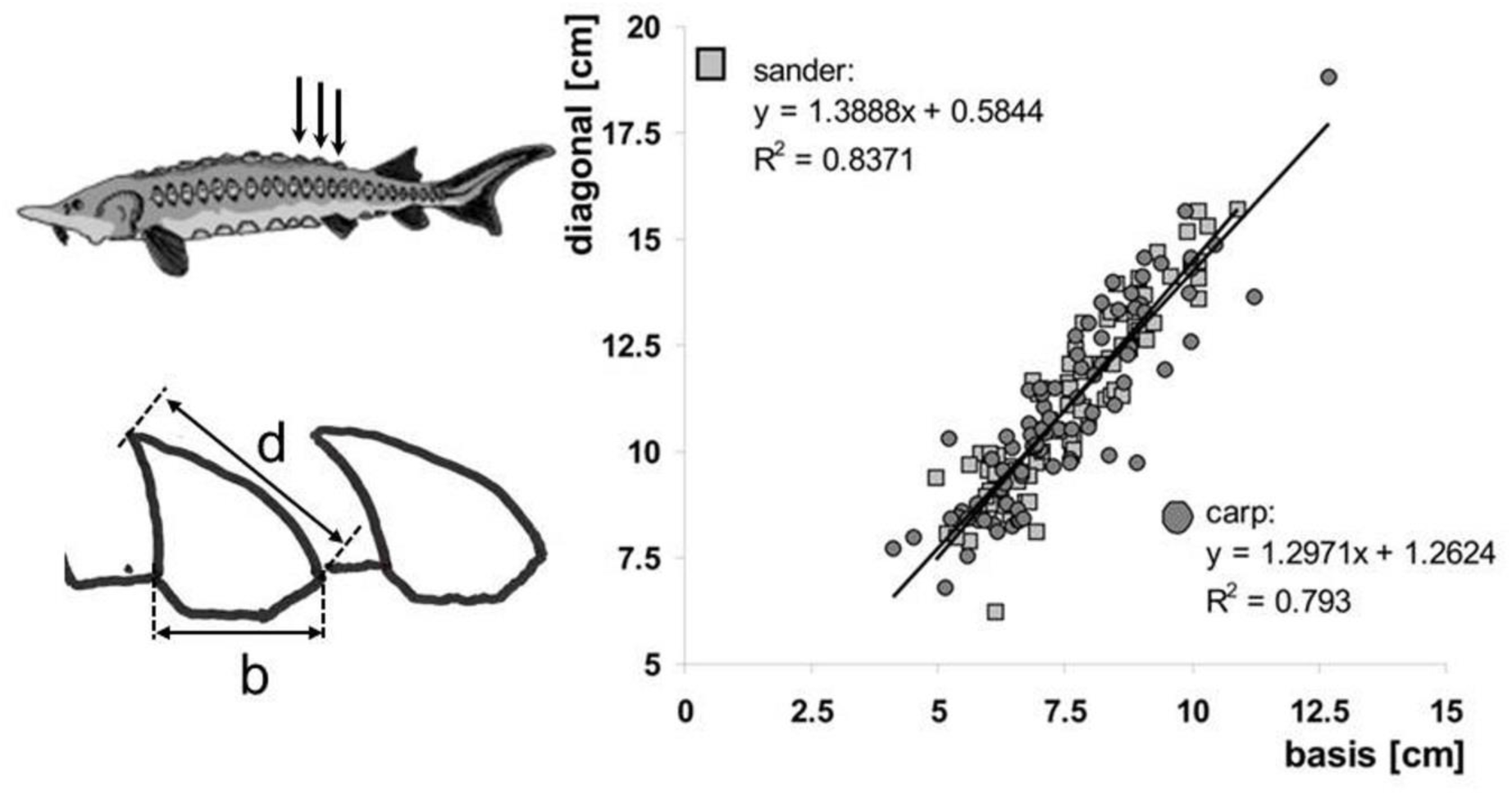
Comparison of the dorsal scutes dimensions (b-basis, d – diagonal) from juvenile Baltic sturgeon *A. oxyrinchus* kept in the rearing water of sander or, as a control, carp for 30 days.

### 3.3. Glucose, lactate and cortisol activities

No significant differences were observed in glucose, lactate or cortisol concentrations in the serum of Baltic sturgeon at the end of the experimental trial (Fig 4). Cortisol concentrations did not reflect any differences between treatments groups.

**Figure 4.**
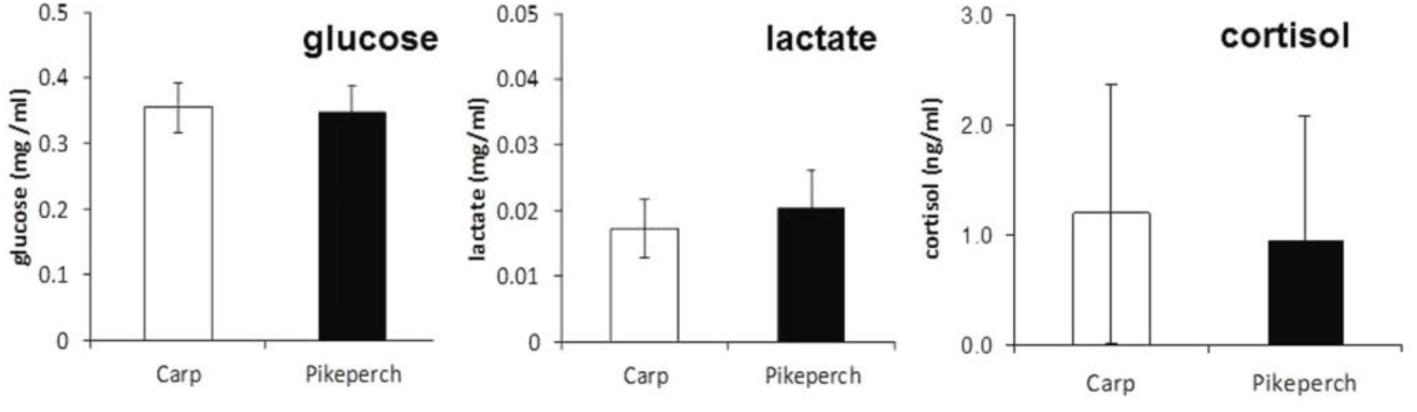
Glucose, lactate and cortisol in the plasma s of juvenile Baltic sturgeon *A. oxyrinchus* kept in the rearing water of sander or, as a control, carp for 30 days. No significant differences between treatments were observed (Mann-Whitney test, p ≤ 0.05).

### 3.4. Brain plasticity and cognition

Selected genes related to brain plasticity and cognition (*neurod1*, *bdnf*, *pcna*) were analyzed in all three brain areas of Baltic sturgeon at the end of the rearing trial. No significant differences were observed in the respective genes for any of the three brain regions (forebrain, midbrain, hindbrain) of Baltic sturgeon (Fig 5).

**Figure 5.**
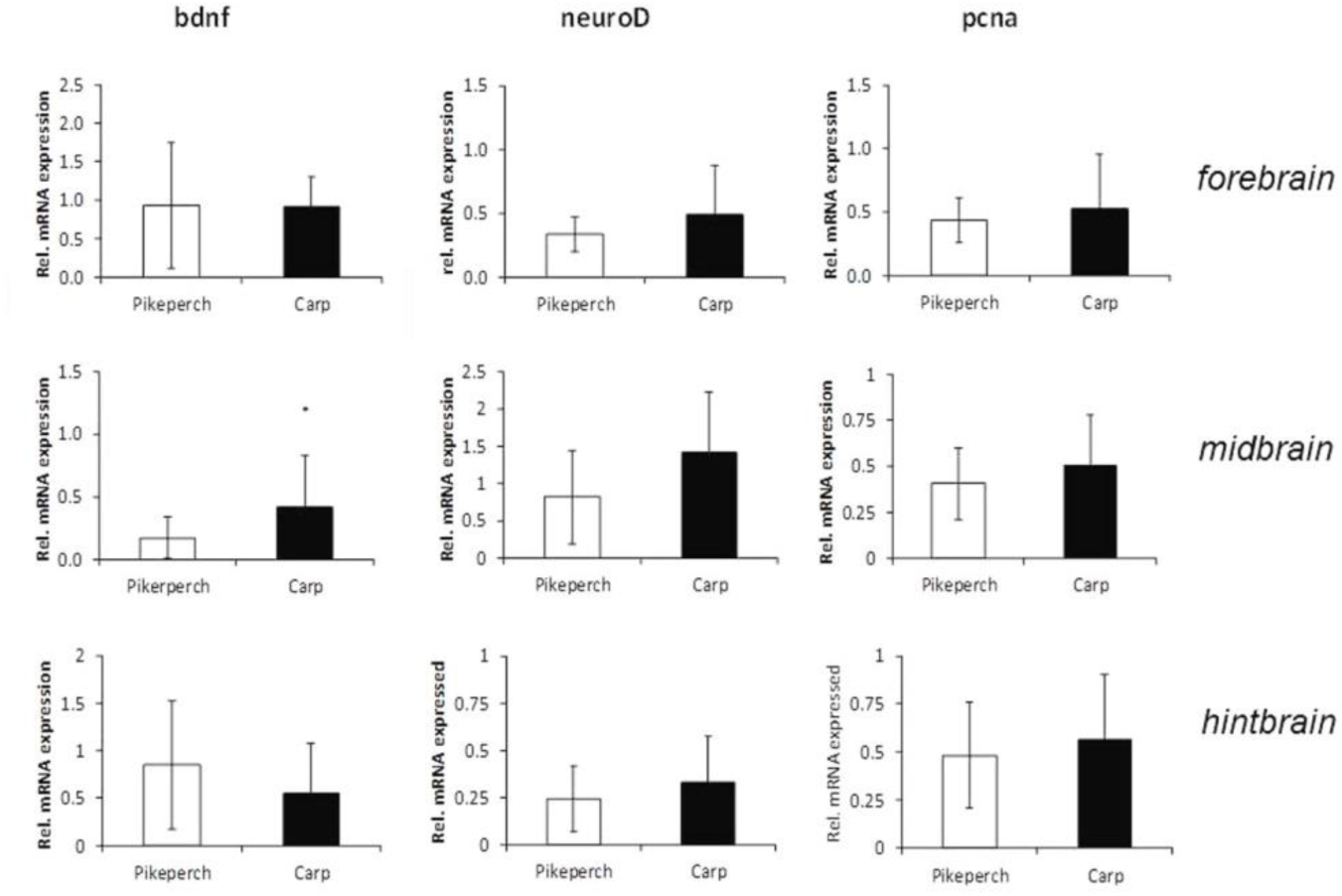
Gene expression of neuroplasticity markers (bdnf, neuroD1, pcna) in the fore-, mid and hintbrain of juvenile Baltic sturgeon *A. oxyrinchus* kept in the rearing water of sander or, as a control, carp for 30 days. No significant differences between treatments were observed (Mann-Whitney test, p ≤ 0.05). Data are expressed as fold relative. bdnf - brain-derived neurotrophic factor (*bdnf*), neurod1 - neurogenic differentiation factor, pcna - proliferating cell nuclear antigen (*pcna*).

## Discussion

Stocking programs routinely release hatchery-reared fish into their natural habitat in an attempt to restore or maintain stable populations. One of the pitfalls of these stocking programs is the elevated mortality of newly stocked individuals (Brown & Smith, 1998; Suboski & Templeton, 1989), which in most cases is highest during and immediately after release (Olla et al.,1994). Unfortunately, methods for reducing post-release mortality have not kept the same pace as aquaculture technology. In general, conservationists have major concerns regarding early life experiences in artificial environments and their impact upon the resulting lack of fitness for survival (Johnsson et al., 2014).

When hatchery-reared fish are released into the wild, they are immediately placed in a novel and variable environment and are also exposed to predatory risk, which they do not experience during the hatchery period. Most mortality during and right after release is due to predation (Brown & Day, 2002) Conservation managers should focus on improving fish survival through reducing mortality for the periods during and immediately following release (Sproul &Tominaga, 1992), which will help in closing the gap between wild and hatchery-reared fish. The differences might be overcome by enriching the environment in which the fish are reared. Furthermore, there is growing evidence that antipredator behaviour is highly sensitive to artificial rearing (Berejikian, 1995). This process should include anti-predator and foraging training before releasing the fish into the wild.

As with other behavior, anti-predator responses have both inherent and learned components. However, most behavior patterns should be viewed as a combination of these two (Kieffer & Colgan, 1992). Learned recognition of novel predators could potentially increase individual’s survival during later predator encounters (Gazdewich & Chivers, 2002; Mirza & Chivers, 2000). Furthermore, several studies have shown that prey reduce their vulnerability to predation by changing morphology, life history strategy and/or behavior when exposed to substances emitted by a predator (Brönmark & Hansson, 2000).

In this study, the objective was to determine if Baltic sturgeon (*A. oxyrinchus*) was able to innately (inherent component) recognize the smell of a common predator, sander (*Sander lucioperca*). In the experiments, there was no indication of differences in the morphology of the dorsal scutes, uneven distribution within the tank, stress levels (glucose, lactate and cortisol) or gene expression of brain plasticity. Thus, we conclude that there was no indication that Baltic sturgeon was chronically stressed during the experimental trial.

Several options should be considered as underlying causes to contributing to this result. First, the fish might not be able to differentiate the smell of the two species when naïve. Second, at the size of the fish tested, carps could be considered a potential predator too. For the behavior observations, the time span between introducing the fish and recording their response has already allowed acclimation. The same reason holds true for the blood stress parameters. In contrast to these hypothetical explanations, the results for the plasticity of the brain regions provide clear evidence that either the control was not functional or that the smell recognition is not effective in naïve fish.

In conclusion, these results suggest that in order to raise Baltic sturgeon with the necessary anti-predator behavior to survive in the wild, it would be necessary to undergo a more complex training period. Only the exposure to predator smell is not sufficient to stimulate anti-predator behavior. In future trials, the training period should consist in implementing pairing predator odour with conspecifics alarm substances. This would be mostly appropriate as it seems that Baltic sturgeon lack anti-predator inherent components. Also, the type of predator exposure could impact antipredator behavior learning. For example, instead of a continuous exposure (ongoing stress), the exposure of Baltic sturgeon to potential predators could take place only intermittently to avoid habituation. It might be needed to be taken into account that as sturgeon develop morphological defenses, such as the growth of pointed bony scutes, they may rely less upon antipredator behavior to prevent being utilized as prey. If morphological or chemical defenses of prey are effective defense mechanisms, it is possible that other behavioral responses may appear to be weak (DeWitt & Langerhans, 2003).

## Acknowledgments

The authors thank Eva Kreuz for her contribution in the laboratory. The study was supported by the European Training Network of the Marie Sklodowska-Curie Actions ITN “Improved production strategies of endangered freshwater species”. This project has received funding from the European Union’s Horizon 2020 research and innovation program under the Marie Sklodowska-Curie grant No [642893].

## Author contributions

The experiment was conducted by A.M. and C.E.S.. The laboratory analysis was carried out by M.C.R. M.C.R. wrote the first draft of the manuscript. S.W. supervised the project. The manuscript was revised by all co-authors.

## References

Berejikian, B.A. (1995). The effects of hatchery and wild ancestry and experience on the relative ability of steelhead trout fry (*Oncorhynchus mykiss*) to avoid a benthic predator. Canadian Journal of Fisheries and Aquatic Sciences, 52, 2476–2482.

Brönmark, C., & Hansson, L.A. (2000). Chemical communication in aquatic systems: An introduction. Oikos, 88, 103–109. https://doi.org/10.1034/j.1600-0706.2000.880112

Brown, C., Day, R.L. (2002). The future of stock enhancements: Lessons for hatchery practice from conservation biology. Fish and Fisheries, 3, 79–94. https://doi.org/10.1046/j.1467-2979.2002.00077

Brown, C., & Laland, K. (2001). Social learning and life skills training for hatchery reared fish. Journal of Fish Biology, 59, 471–493. https://doi.org/10.1006/jfbi.2001.1689

Brown, G.E., & Smith, R.J.F. (1998). Acquired predator recognition in juvenile rainbow trout (*Oncorhynchus mykiss*): conditioning hatchery-reared fish to recognize chemical cues of a predator. Canadian Journal of Fisheries and Aquatic Sciences, 55, 611–617. https://doi.org/10.1139/f97-261

DeWitt, T.J., & Langerhans, R.B. (2003). Multiple prey traits, multiple predators: Keys to understanding complex community dynamics. Journal of Sea Research, 49, 143–155. https://doi.org/10.1016/S1385-1101(02)00220-4

Ferno, A., & Jarvi, T. (1998). Domestication genetically alters the antipredator behaviour of anadromous brown trout (*Salmo trutta*)- dummy predator experiment. Nordic Journal of Freshwater Research, 74, 95–100.

Ferrari, M.C.O., Wisenden, B.D., & Chivers, D.P. (2010). Chemical ecology of predator–prey interactions in aquatic ecosystems: a review and prospectus‥ Canadian Journal of Zoology, 88, 698–724. https://doi.org/10.1139/Z10-029

Gazdewich, K.J., & Chivers, D.P. (2002). Acquired predator recognition by fathead minnows: Influence of habitat characteristics on survival. Journal of Chemical Ecology, 28, 439–445.

Gessner, J., Arndt, G.M., Tiedemann, R., Bartel, R., & Kirschbaum, F. (2006). Remediation measures for the Baltic sturgeon: Status review and perspectives. Journal of Applied Ichthyology, 22, 23–31. https://doi.org/10.1111/j.1439-0426.2007.00925

Gessner, J., Tautenhahn, M., Spratte, S., Arndt, G.M., & von Nordheim, H. (2011). Development of a German Action Plan for the restoration of the European sturgeon *Acipenser sturio* L. - implementing international commitments on a national scale. Journal of Applied Ichthyology, 27, 192–198. https://doi.org/10.1111/j.1439-0426.2011.01697

Griffin, A.S., Blumstein, D.T., & Evans, C.S. (2000). Training captive-bred or translocated animals to avoid predators. Conservation Biology, 14, 1317–1326. https://doi.org/10.1046/j.1523-1739.2000.99326

IUCN (2018). The IUCN Red List of Threatened Species. Version 2018-1.

Johnsson, J.I., & Abrahams, M.V. (1991). Interbreeding with domestic strain increases foraging under threat of predation in juvenile steelhead trout (*Oncorhynchus mykiss*): An experimental study. Canadian Journal of Fisheries and Aquatic Sciences, 48, 243–247. https://doi.org/https://doi.org/10.1139/f91-033

Johnsson, J.I., Brockmark, S., & Näslund, J. (2014). Environmental effects on behavioural development consequences for fitness of captive-reared fishes in the wild. Journal of Fish Biology, 85, 1946–1971. https://doi.org/10.1111/jfb.12547

Kelley, J.L., & Magurran, A.E. (2003). Learned predator recognition and antipredator responses in fishes. Fish and Fisheries, 4, 216–226. https://doi.org/10.1046/j.1467-2979.2003.00126

Kieffer, J.D. & Colgan, P. (1992). The role of learning in fish behavior. Reviews in Fish Biology and Fisheries, 2, 125–43.

Langhans, S.D., Gessner, J., Hermoso, V., & Wolter, C. (2016). Coupling systematic planning and expert judgement enhances the efficiency of river restoration. Science of the Total Environment, 560–561, 266–273.https://doi.org/10.1016/j.scitotenv.2016.03.232

Mirza, R.S., & Chivers, D.P. (2000). Predator-recognition training enhances survival of brook trout: evidence from laboratory and field-enclosure studies. Canadian Journal of Zoology, 78, 2198–2208. https://doi.org/10.1139/z00-164

Olla, B.L., Davis, M.W., & Ryer, C.H. (1998). Understanding how the hatchery environment represses or promotes the development of behavioral survival skills. Bulletin of Marine Sciences, 62, 531–550.

Olla, B.L., Davis, M.W. & Ryer, C.H. (1994). Behavioural deficits in hatchery-reared fished: Potential effects on survival following release. Aquaculture and Fisheries Management, 25, 19–34.

Pfaffl, M.W. (2001). A new mathematical model for relative quantification in real-time RT-PCR. Nucleic Acids Research, 29, 45.

Reiser, S., Wuertz, S., Schroeder, J.P., Kloas, W., Hanel, R. (2011). Risks of seawater ozonation in recirculation aquaculture - Effects of oxidative stress on animal welfare of juvenile turbot (*Psetta maxima*, L.). Aquatic Toxicology, 105, 508–517. https://doi.org/10.1016/j.aquatox.2011.08.004

Sproul, T., & Tominaga, O. (1992). An economic review of the Japanese flounder stock enhancement project in Ishikary Bay. Bulletin of Marine Science, 50, 75–88.

Suboski, M.D., & Templeton, J.J. (1989). Life skills training for hatchery fish: Social learning and survival. Fisheries Research, 7, 343–352. https://doi.org/10.1016/0165-7836(89)90066-0

Wallace, M.P. (2000). Retaining natural behavior in captivity for reintroduction programmes, In Behaviour and Conservation (eds L.M. Gosling and W.J. Sutherland). Cambridge University Press, Cambridge, pp. 300–314.

